# Mitonuclear sex determination? Empirical evidence from bivalves

**DOI:** 10.1101/2023.07.05.547839

**Authors:** Chase H. Smith, Raquel Mejia-Trujillo, Sophie Breton, Brendan J. Pinto, Mark Kirkpatrick, Justin C. Havird

**Affiliations:** Department of Integrative Biology, University of Texas at Austin, Austin, TX, USA; Department of Biological Sciences, University of Montreal, Montreal, Canada; School of Life Sciences, Arizona State University, Tempe, AZ USA; Center for Evolution and Medicine, Arizona State University, Tempe, AZ USA; Department of Zoology, Milwaukee Public Museum, Milwaukee, WI USA

**Keywords:** Sexual development, mitonuclear interactions, doubly uniparental inheritance, Unionida, mt-sncRNAs

## Abstract

Genetic elements encoded in nuclear DNA determine the sex of an individual in many animals. In bivalves, however, mitochondrial DNA (mtDNA) has been hypothesized to contribute to sex determination in lineages that possess doubly uniparental inheritance (DUI). In these cases, females transmit a female mtDNA (F mtDNA) to all offspring, while male mtDNA (M mtDNA) is transmitted only from fathers to sons. Because M mtDNA is inherited in the same way as Y chromosomes, it has been hypothesized that mtDNA may be responsible for sex determination. However, the role of mitochondrial and nuclear genes in sex determination has yet to be validated in DUI bivalves. In this study, we used DNA, RNA, and mitochondrial short non-coding RNA (sncRNA) sequencing to explore the role of mitochondrial and nuclear elements in the sexual development pathway of the freshwater mussel *Potamilus streckersoni* (Bivalvia: Unionida). We found that the M mtDNA shed a sncRNA partially within a male-specific mitochondrial gene that targeted pathways hypothesized to be involved in female development and mitophagy. RNA-seq confirmed the gene target was significantly upregulated in females, supporting a direct role of mitochondrial sncRNAs in gene silencing. These findings support the hypothesis that M mtDNA inhibits female development. Genome-wide patterns of genetic differentiation and heterozygosity did not support a nuclear sex determining region, although we cannot reject that nuclear factors are involved with sex determination. Our results provide further evidence that mitochondrial loci contribute to diverse, non-respiratory functions and provide a first glimpse into an unorthodox sex determining system.

## Introduction

In many animals, sex is determined by genetic elements encoded in nuclear DNA, and mitochondrial DNA (mtDNA) has not been demonstrated to play a role in sex determination (Bachtrog et al. 2014). However, mechanisms underlying sex determination are highly variable across the tree of life and have been shown to include both mitochondrial and nuclear gene products in plants (Hanson and Bentolila 2004). mtDNA has also been hypothesized to play a role in sex determination or sexual development in bivalves that possess a unique mitochondrial biology known as doubly uniparental inheritance (DUI) (Breton et al. 2011; Breton et al. 2022).

Sex determining pathways in bivalves with DUI have been of interest to researchers since its discovery in the early 1990’s (Hoeh et al. 1991). Doubly uniparental inheritance involves the biparental transmission of mtDNA, one passed by females to all offspring and a second transmitted by males to only male offspring (Skibinski et al. 1994) (Fig. 1). Females typically only possess female-transmitted mtDNA (F mtDNA), while males are often globally heteroplasmic in somatic tissues but male-transmitted mtDNA (M mtDNA) is localized in gonads and exclusively possessed in sperm (Breton et al. 2017; Ghiselli et al. 2019). Given M mtDNA are inherited in the same way as Y chromosomes and associated with maleness, M mtDNA has been hypothesized to trigger development of male phenotypes, which is hypothesized to be suppressed when M mtDNA is degraded and only F mtDNA remains (Breton et al., 2007).

**Figure 1.**
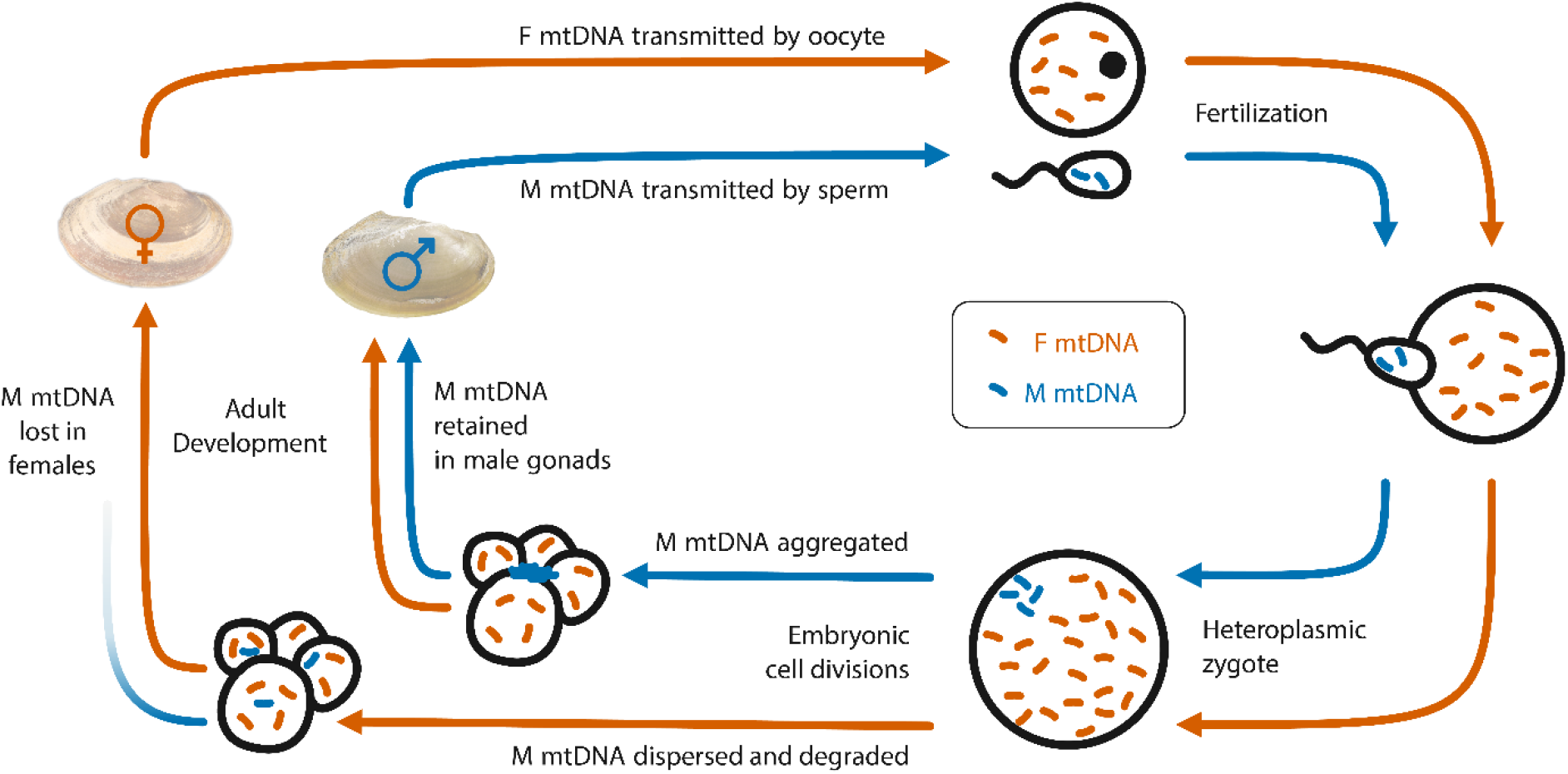
Overview of doubly uniparental inheritance of mitochondria in bivalves. Orange coloration represents the female-transmitted (F) mitochondrial DNA (mtDNA) and blue male-transmitted (M) mtDNA. Adapted from Breton et al. (2018).

In freshwater mussels (Bivalvia: Unionida), the evolutionary conservation of DUI and unique sex-specific mitochondrial genes has led to the hypothesis that mtDNAs could play a direct role in sex determination (Breton et al. 2011; Breton et al. 2022). Unlike most other DUI lineages where F and M mtDNA divergence is often ∼15% or less, the F and M mtDNAs of freshwater mussels are more than 50% divergent in their amino acid sequences and have remained evolutionarily distinct for over 200 million years (Breton et al. 2007; Doucet-Beaupré et al. 2010; Smith et al. 2023). Further, both genomes contain open reading frames (ORFs; termed female ORF [F-ORF] and male ORF [M-ORF]) with no known homology that have been hypothesized to be either primary sex determining gene(s) or contribute to sexual development (Breton et al. 2011). This mtDNA sex determination hypothesis has been further supported by evolutionary transitions from dioecy to hermaphroditism leading to divergent evolution in F-ORF peptide sequence of hermaphrodites relative to other mitochondrially encoded proteins (Breton et al. 2011; Mitchell et al. 2016; Guerra et al. 2019). Because of these characteristics, freshwater mussels have become an ideal model for investigating mitochondrial biology broadly and sex determination hypotheses for DUI species and bivalves in general.

We highlight two specific hypotheses regarding the retention of M mtDNA and the role of mtDNAs in the sexual development of DUI species: (1) In the egg factor hypothesis, sex is determined by an egg factor contributed by the mother (Kenchington et al. 2009; Ghiselli et al. 2012; Milani et al. 2013; Zouros and Rodakis 2019; Zouros 2020). The egg factor is an inherited genotype at a nuclear gene (or possibly multiple genes) that triggers sexual development, including the retention of the M mtDNA and its localization in male gonadal cells. This hypothesis generally follows expectations under a traditional XY or ZW sex determination system and does not account for the potential role that mitochondrial-encoded elements may play in sexual development. (2) In the cytoplasmic male sterility (CMS) hypothesis, which is primarily based on evidence from freshwater mussels, mtDNAs play a direct role in sexual development (Breton et al. 2022). In the model, the F-ORF acts as a feminizer (inhibiting maleness) and the M-ORF antagonizes F-ORF in some way, acting as a ‘restorer of maleness’, leading to downstream interactions with nuclear gene products to trigger development of male phenotypes. These two hypotheses provide a foundation for future research to test if one or both hypothetical pathways are involved in sexual development in DUI bivalves.

The CMS hypothesis provides an explanation for the evolutionary conservation of F-ORF and M-ORF across freshwater mussels, but it remains uncertain whether these genes encode proteins that trigger sexual development, if they interact with nuclear gene products, or if other regulatory elements encoded in mtDNAs interact with the nuclear genome. In the nuclear genome, short non-coding RNAs (sncRNAs) are critical factors that regulate nuclear gene expression typically through RNA interference (RNAi), a process in which sncRNA binding blocks translation of target messenger RNA (Ambros 2004). sncRNAs shed by mtDNA (mt-sncRNAs) have been identified and confirmed to alter nuclear gene expression (Pozzi and Dowling 2021; Pozzi and Dowling 2022), including in DUI species (Pozzi et al. 2017; Passamonti et al. 2020). However, mt-sncRNA validation has yet to be performed in freshwater mussels and their contribution to sexual development, therefore, remains uncertain.

In this study, we set out to explore the role of mitochondrial and nuclear elements in the sexual development of the freshwater mussel *Potamilus streckersoni*. Specifically, we use whole genome resequencing, female and male gonad RNA sequencing, and female and male gonad mt-sncRNA sequencing in attempts to identify a candidate SDR within the nuclear genome and mitochondrial-encoded factors contributing to sexual development. We then synthesize our results with previous sex determination hypotheses presented in the literature. Our findings provide further support for an unusual sex determination system in DUI bivalves dependent on mitonuclear interactions.

## Materials and Methods

### DNA extraction and sequencing

To investigate nuclear genes that may be involved with sex determination, we collected 22 adult individuals of *P. streckersoni* from multiple localities in the Brazos River drainage in Texas. This population has been shown to lack population structure based on genomic data (Smith et al. 2021). Sex was distinguished using shell characters (adult females typically have blunt posterior shell margins) and external gill morphology (adult females have distinct serrated gill margins). DNA was extracted from fresh mantle tissue using the Qiagen PureGene Kit (Hilden, Germany) with standard protocols. High-molecular weight genomic DNA was confirmed from these extractions by visualizing each isolation on a 1% agarose gel stained with GelGreen® nucleic acid stain (Biotium, Hayward, CA, USA). Isolation quantity and quality was assessed using a Qubit™ fluorometer and a NanoDrop™ One (ThermoFisher Scientific; Waltham, MA, USA), respectively.

Whole genome resequencing (WGR) was performed on 11 female and 11 male individuals of *P. streckersoni*. Libraries were prepared by the Texas A&M AgriLife Genomics and Bioinformatics Service (College Station, TX, USA) from ∼25 ng of genomic DNA in a custom, automated, and miniaturized version of the PerkinElmer NEXTFLEX Rapid XP kit protocol (Johnson et al. 2019). Briefly, genomic DNA was enzymatically fragmented for 6 minutes, ligated to unique dual-indexed barcodes, size selected between 520–720 bp using SPRIselect beads (Beckman Coulter; Brea, CA, USA), and amplified with 10 PCR cycles. One tenth of the manufacturer’s prescribed volumes were used for enzymatic steps. Libraries were diluted with elution buffer to a final concentration of 2 ng/μl and were pooled by equivolume. The pool was sequenced on an Illumina NovaSeq S4 XP (San Diego, CA, USA) using 2 × 150 bp reads.

### DNA Analyses

We used pool and individual based single nucleotide polymorphism (SNP) approaches to further investigate a potential nuclear SDR in *P. streckersoni*. Raw reads were trimmed using TRIM GALORE! V 0.6.7 (www.bioinformatics.babraham.ac.uk/projects/trim_galore/) with default settings and data quality was verified in FastQC v 0.11.9 (www.bioinformatics.babraham.ac.uk/projects/fastqc/). Trimmed reads were mapped to the *P. streckersoni* nuclear genome assembly (Smith 2021) using bwa-mem v 2.2.1 (Md et al. 2019) with default parameters. PCR duplicates were removed using picard v 2.27.4 (Broad Institute 2019). The resulting bam files were combined by sex and the Pooled Sequencing Analyses for Sex Signal (PSASS) pipeline v 3.1.0 (Feron and Jaron 2021) was used to call SNPs and calculate genotypic statistics. Pileup was used to call SNPs with a minimum mapping quality of 30. We calculated sex-specific alleles within gene models using PSASS. We then calculated F_ST_ between the sexes using 1 kb windows and considered windows with values greater than or equal to 0.1 as elevated. Windows with at least an average of 3x coverage per individual per sex (33x) were retained for the analysis.

Pool-based methods failed to identify large, contiguous areas of differentiation between the sexes. To identify smaller regions of the genome with alleles consistent with XY or ZW sex determination (XY- or ZW-like alleles), we investigated patterns of heterozygosity, which have been used previously to identify SDRs in some bivalves (Han et al. 2022). First, we downloaded high coverage Illumina reads for the individual used to generate the reference assembly from the GenBank SRA (accession SRR13176629). We then recalled SNPs including all 22 WGR samples and the reference individual using FreeBayes v 1.3.6 (Garrison and Marth 2012). Variants were filtered to only include Q20 biallelic sites, and singletons were removed using vcftools v 0.1.16 (Danecek et al. 2011). We then followed similar methods as Kirkpatrick et al. (2022) to identify ZW or XY-like alleles. For ZW-like SNPs, the script required all males to be homozygous for the same allele and females heterozygous with one copy of the same allele found in males. For XY-like SNPs, we used the inverse. Although SNPs following this pattern could occur across the genome by chance, they are expected to be enriched in SDRs. For our analysis, we required called genotypes from at least 3 males and 3 females at each candidate site. We used a modified version of the R script provided in Kirkpatrick et al. (2022) to perform the analysis.

### RNA extraction and sequencing

We extracted total RNA from fresh or preserved (RNAlater) gonadal tissue of 6 female and 3 male *P. streckersoni* to infer differential gene expression among nuclear genes. To improve the existing genome annotation (Smith 2021), we also extracted RNA from six tissue types from an adult male *P. streckersoni*: adductor, foot, gill, gonad, mantle, and stomach. RNA was extracted from each tissue type independently and pooled into a single sample with equal representation from RNA samples. All RNA was extracted using the RNeasy kit following the manufacturer’s protocol (Qiagen). RNA quality and integrity were determined using a NanoDrop^TM^ and an Agilent Bioanalyzer (Santa Clara, CA, USA), respectively. Messenger RNA was isolated from 150 ng of total RNA using a Nextflex Poly-A Selection kit (Perkin Elmer; Waltham, MA, USA). cDNA libraries were prepared using a Nextflex Rapid Directional RNA 2.0 kit, miniaturized to 2/5 reaction volume and automated on a Sciclone NGSx liquid handler. All libraries were sequenced by the Texas A&M AgriLife Genomics and Bioinformatics Service on an Illumina NovaSeq S4 XP using 2 × 150 bp reads.

### Potamilus streckersoni *genome annotation*

We used all novel RNA-Seq reads, as well as female and larval pools published previously (SRA accession numbers SRR13176627 and SRR13176628), to improve the existing genome annotation for *P. streckersoni* (Smith 2021). Structural and function annotation was performed using the Funannotate pipeline v 1.8.10 (Palmer and Stajich 2017). Prior to running the pipeline, repeats in the *P. streckersoni* genome assembly were identified and masked using RepeatModeler v 2.0.1 (Flynn et al. 2020) and RepeatMasker v 4.0.9 (Smit et al. 2015), respectively. Completeness of the previous and updated annotation was assessed using BUSCO v 5.4.3 (Manni et al. 2021) with the metazoan (v 10; 954 genes) and molluscan lineages (v 10; 5295 genes).

### RNA-seq analyses

Gonadal RNA-Seq reads were trimmed using TRIM GALORE! with default parameters. Data quality was verified using FastQC. Trimmed reads were then used to test for significant differences in gene expression between sexes using the updated genome annotation. Reads were mapped to the reference genome using Hisat2 v 2.2.1 (Kim et al. 2019) with default parameters. Gene counts were summarized using the command “featureCounts” in the R package Rsubread v 2.10.5 (Liao et al. 2019). A differential gene expression analysis between males and females was performed on the counts using the command “DESeq” in the R package DESeq2 v 1.36.0 (Love et al. 2014) with default parameters. Genes with adjusted p-values < 0.05 were considered differentially expressed genes (DEGs) among sexes. We then joined expression data for genes that fell within windows of high F_ST_ based on DNA analyses and tested whether genes that fell within windows of high F_ST_ were more likely to be differentially expressed using a chi-squared test.

To infer co-expression among genes, we performed a weighted gene co-expression network analysis (WGCNA) in the R package WGCNA v.1.72-1 (Langfelder and Horvath 2008). Before the analysis, we filtered out genes with a coefficient of variation > 200 and counts per million value < 5 from the raw count matrix using the “filtered.data” command in the R package NOISeq v 2.40.0 (Tarazona et al. 2015). The count data was then transformed using the “vst” command in DESeq2. Once the co-expression network was constructed following developers’ documentation, we used eigengenes as representatives of each module to investigate intermodular correlations. Additionally, we included binary traits (female and male) to the eigengene network to reveal co-expression relationships across the sexes. We identified genes with high intramodular connectivity (*i.e.*, hub genes) using the “intramodularConnectivity” function from the WGCNA package.

We further investigated potential pathways that were differentially expressed among female and male gonads using pre-ranked gene set enrichment analyses in the R package fgsea v 1.22.0 (Korotkevich et al. 2021). We used the hallmark (Liberzon et al. 2015) and GO Biological Process gene sets from the Human Molecular Signatures Database v 2022.1.Hs (MSigDB) (Subramanian et al. 2005) for the analyses. Gene names from functional annotations from funannotate were edited to match human gene symbols from entries in MSigDB, while genes without annotations were removed. Z-statistics from DESeq2 were used to rank genes. Additionally, the Z-scores of duplicated genes were averaged such that each unique gene symbol had one value contributing towards its rank. Pre-ranked gene set enrichment analyses were then performed on each gene set using the command “fgseaMultilevel” in fgsea with a maximal gene set size of 50. We considered pathways with an adjusted p-value < 0.05 as having significantly different expression among female and male gonadal tissue.

### Mitochondrial sncRNA sequencing, validation, and target prediction

High throughput sncRNA-seq was performed on gonadal tissue from 3 females and 3 males, which included both new individuals and a subset of individuals used for RNA-Seq and DEG analyses. Before sncRNA extraction, mitochondrial enrichment was performed on gonadal tissue samples using a modified protocol described in Ballantyne & Moon (1985). Briefly, this process involved an initial homogenization in ice-cold isolation buffer (480 mM sucrose, 100 mM KCl, 50 mM NaCl, 70 mM HEPES, 6 mM EGTA, 3 mM EDTA, and 1% BSA; pH 7.6) followed by two centrifugation steps: 1) homogenates centrifuged at 4°C at 2,790 X g for 5 minutes, and 2) supernatants centrifuged 4°C at 12,200 X g for 15 minutes. We used the Purelink miRNA extraction kit (Invitrogen) with the standard protocol to extract sncRNA from pellets enriched with mitochondria. Libraries were prepared for each sample at the University of Texas Genomic Sequencing Facility using the NEBNext Small RNA library preparation kit (New England Biolabs; Ipswich, MA, USA). Libraries were enriched with 14 cycles of PCR and the final product was size selected for 105–155 bp using a 3% gel cassette on the Blue Pippin (Sage Science; Beverly, MA, USA). The final size selected libraries were assessed for library quality and quantity using a BioAnalyzer. Libraries were subsequently sequenced on an Illumina NovaSeq 6000 using 1 × 75 or 1 × 100 bp reads.

Data processing generally followed previous studies identifying mt-sncRNAs (Pozzi et al. 2017; Passamonti et al. 2020). Raw reads were trimmed using TRIM GALORE! while enforcing a minimum read length of 15 bp and a maximum of 40 bp. Trimmed reads were then mapped to reference female (GenBank: MW413895) and male (GenBank: ON881148) mitochondrial genome assemblies for *P. streckersoni* (Smith 2021; Mejia-Trujillo and Smith 2022) using bowtie v 2.5.0 (Langmead and Salzberg 2012) with the parameters “-N 1-i C,1-L 18--end-to-end”. Mapped reads were then clustered using the UCLUST algorithm in USEARCH V 11.0.667 (Edgar 2010) using an identity filter of 0.99. Clusters with centroids less than 17 bp were removed. Centroids that followed the following criteria were considered mitochondrial sncRNAs: (1) a cluster size greater than 200; (2) a perfect match of nucleotides 4–10 and a minimum of 11 matches with a 3’ UTR of a nuclear-encoded gene as determined by BLAST + 2.6.0105 (Camacho et al. 2009) using the options “-task blastn-short-strand minus”; and (3) a ΔΔG score lower than −9 kJ for the centroid-target UTR interaction as determined by RNAup v 2.5.1 (Lorenz et al. 2011) using a temperature of 37 °C, (4) a Gibbs free energy score lower than −20 kJ for the centroid-mRNA duplex as determined by RNAhybrid v 2.1.2 (Krüger and Rehmsmeier 2006), and (5) at least a 1.5 fold difference in coverage at the 5’ and 3’ ends when compared to the average value for 5 bp upstream and downstream as determined by SAMtools v 1.6 (Danecek et al. 2021). In cases where targets were annotated as hypothetical proteins, we annotated genes (if able) by blasting the peptide sequence against the Kyoto Encyclopedia of Genes and Genomes (KEGG) (Kanehisa et al. 2023). We then examined expression data for mt-sncRNA targets and tested whether targeted genes were significantly more likely to be differentially expressed than those that were not using a chi-squared test.

### Gene network analysis and protein-protein interaction prediction

Our results identified a particularly interesting M mt-sncRNA at the 5’ end of the M-ORF targeting the nuclear-encoded gene *GCNT1*, prompting us to use STRING v 11.5 (Szklarczyk et al. 2019) to investigate the gene interaction network of *GCNT1*. For the network, we only considered validated interactions based on the following settings in STRING: 1) supported by STRING databases, 2) supported by experimental evidence, 3) and predicted interactions with a confidence score ≥ 0.15. We then visualized gene expression among genes in the network based on expression data from DESeq2.

To infer if the F-ORF protein might interact with the *GCNT1* protein, we used AlphaPulldown v 0.22.3 (Yu et al. 2023) and AlphaFold v 2.2.0 (Jumper et al. 2021). This tested the hypothesis that the F-ORF protein is acting as a feminizer by interaction with the *GCNT1* protein, while the M mt-sncRNA is acting as a restorer of maleness through RNAi-mediated gene silencing. To control for random protein-protein interaction (PPI) support due to the small size of the F-ORF protein (88 residues), we ran the same analysis on the F mtDNA copy of ATP8 (72 residues) and the M-ORF (214 residues). Our logic behind this is that short peptide sequences or mitochondrially based proteins would show similar support for PPI if a supported F-ORF interaction was due to chance. We considered a protein docking score greater than or equal to 0.23 as a predicted PPI (Basu and Wallner 2016).

## Results

### No support for nuclear encoded SDR

Whole genome resequencing performed on 11 male and 11 female individuals of *P. streckersoni* generated approximately ∼7x coverage per sample. Additional statistics regarding samples, sequencing, voucher numbers, and accession numbers can be found in Table S1. Sliding window statistics failed to identify large, contiguous areas of sequence or read depth differentiation between the sexes (File S1), which would be expected if sex chromosomes or large, continuous regions were responsible for sex determination. However, we identified 5629 1-kb windows with elevated F_ST_ (0.1) across 880 scaffolds (roughly ⅓ of total scaffolds) (File S2). This included 2439 windows falling within 1346 genes (File S2). Genome-wide patterns of heterozygosity did not identify any regions that were enriched with XY- or ZW-like SNPs, with only two ZW-like SNPs (in two genes on two different scaffolds) and one XY-like SNPs (not within a gene) identified by our analysis. All SNPs were only present in areas of low coverage (no more than five individuals of either sex) and coinciding sliding windows showed no evidence of high genetic differentiation, which would be expected if the region was a SDR. Therefore, we consider that these SNPs occurred by chance and are most likely not SDRs.

### Improved genome annotation and differential expression between the sexes

Statistics regarding RNA-seq libraries can be found in Table S2. Functional annotation returned 45,268 gene models (34,937 protein-coding genes, 2,075 isoforms, 8,256 tRNAs), reducing the number of protein coding genes from the previous annotation by ∼15%. Despite the decreased number of protein coding genes, the annotation had more than a 20% increase in complete BUSCOs using the metazoan and molluscan lineages (up to 95% and 86% complete, respectively, Table S3).

DESeq2 identified 2311 DEGs (adjusted p-value < 0.05) between female and male gonadal tissue (Fig. 2A; File S3). Of these DEGs, 141 were also found to have at least 1 window of elevated F_ST_ (0.1) (File S2). A chi-squared test supported the hypothesis that genes with elevated F_ST_ were significantly more likely to be differentially expressed (6.1% vs 3.7%; p < 0.001), which is coincident with sexual antagonism in female and male gonadal tissues.

**Figure 2.**
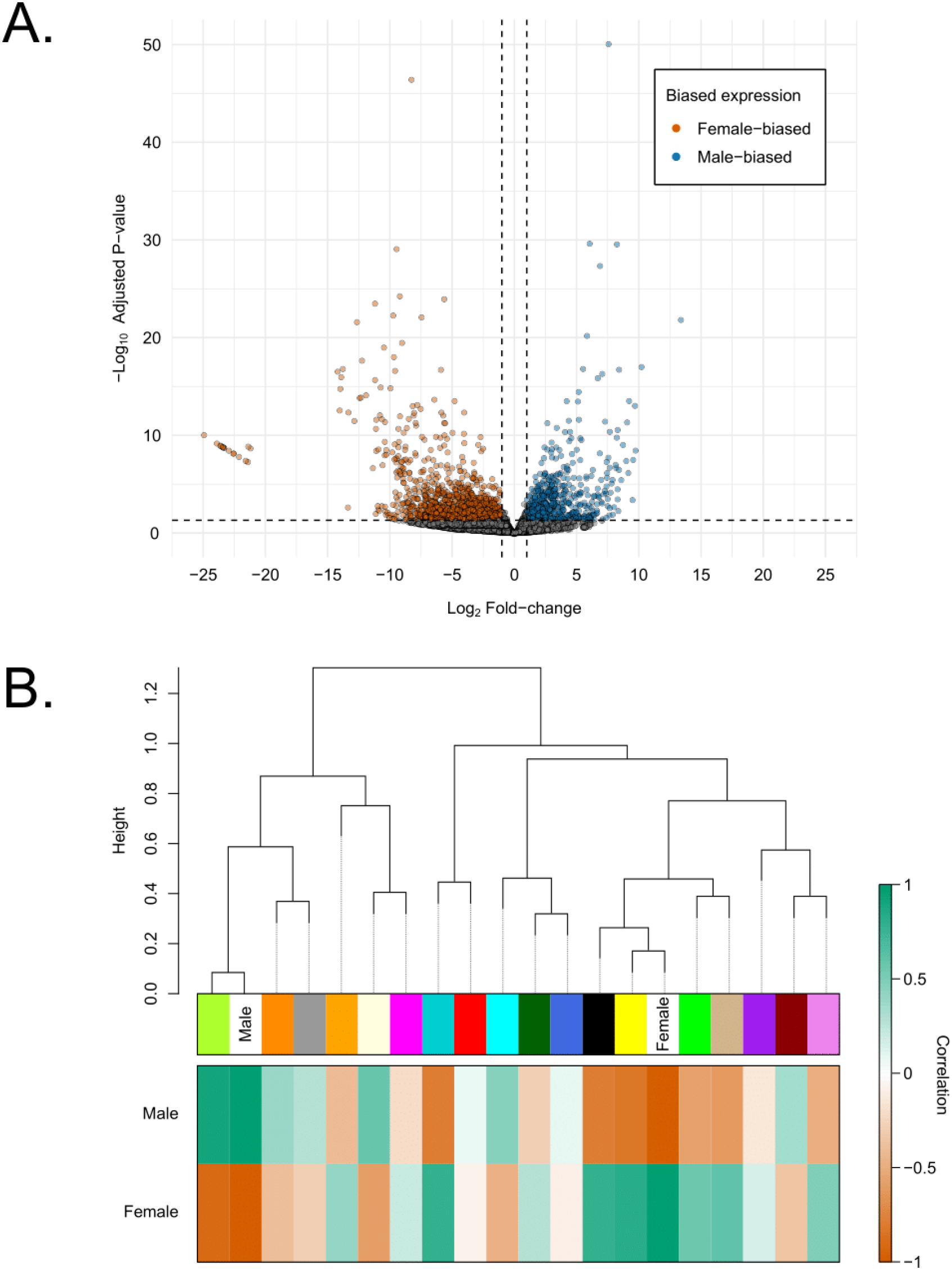
A) Volcano plot summarizing tests for differential gene expression between female and male gonadal tissue in *Potamilus streckersoni*. Orange dots represent genes found to have significant female-biased expression and blue those with significant male-biased expression. Black dots represent genes without significantly different expression. Dashed vertical lines represent a log_2_-fold change of -1 and 1, and dashed horizontal lines represent the - log_10_ value for adjusted p-value of 0.05. B) Dendrogram of WGCNA clusters based on expression profiles. In the heat plot, coloration represents positive (green) or negative (orange) correlation of each cluster with sex (female or male).

WGCNA selected 18 modules to best explain expression profiles (Fig. 2B; File S4). The genes with the highest intramodular connectivity for each module (*i.e.*, the gene with the highest level of co-expression with other genes in the module) are reported in File S5. Genes in the top 5% of intramodular connectivity were found in three modules: 1) 452 of the 1,620 genes in the orange module, 2), 85 of the 1,140 genes in the violet module, and 3) 49 of the 1,909 genes in the green-yellow module (most closely corresponded to maleness). The yellow module consisted of nearly ⅓ of all annotated genes (11,725 genes) and most closely corresponded to femaleness. Additionally, 13 pathways were supported as enriched in the hallmark set and 246 pathways in the GOBP set (p < 0.05) (Fig. 3; File S6). Many of these pathways were involved with sex maintenance and not primary sex determination (*e.g.*, spermatogenesis gene enrichment in males).

**Figure 3.**
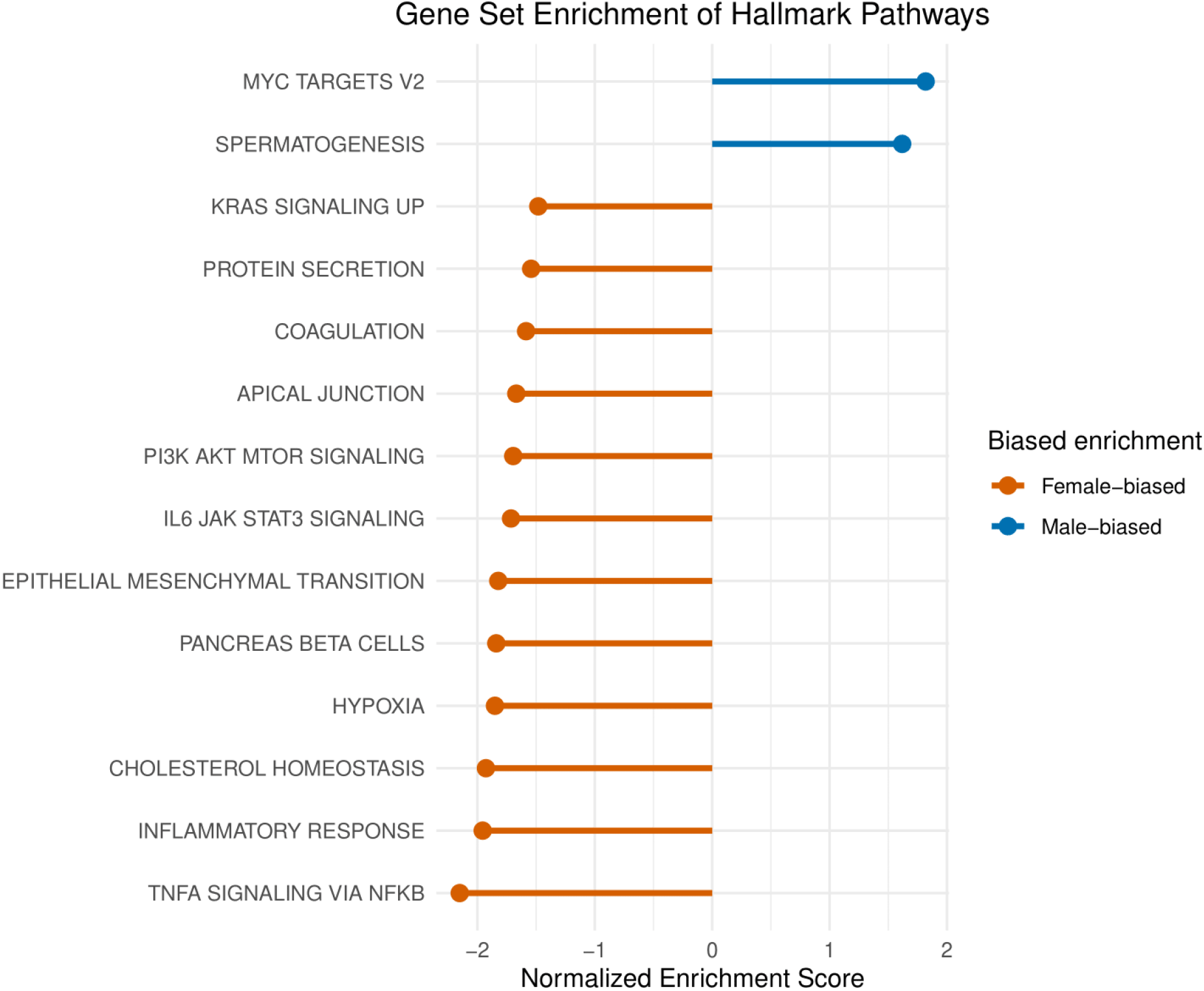
Gene set enrichment of Hallmark Pathways. Pathways with negative NES values contain genes with female-biased expression, while pathways with positive NES values contain genes with male-biased expression. All shown pathways showed significant enrichment (p-adj < 0.05) in female (orange) and male (blue) gonadal tissue.

### Female and male mitochondrial sncRNAs inhibit nuclear gene expression

High throughput sncRNA-seq generated an average of 32 million reads per sample (female: 37 million, male: 29 million). Additional details regarding each library can be found in Table S4. In total, 24 F and nine M mt-sncRNAs passed our inclusion criteria. All mt-sncRNAs, their location, and validated targets are reported in Table S5. In total, 91 nuclear targets were identified for the 34 mt-sncRNAs. The mt-sncRNAs were highly enriched, with read counts that are orders of magnitude greater than in mt-sncRNAs previously described from DUI bivalves (Pozzi et al., 2017). We noticed one male sample expressed F mt-sncRNAs at a higher level than the other male samples (Fig. 4). However, this is reasonable and likely due to chance given our sampling design, because male gonadal tissue could include cells containing F mtDNA (only sperm show homoplasmy for M mtDNA) and M mtDNA concentration in gonadal tissue can vary based on individual or seasonality in *P. streckersoni* (Mejia-Trujillo and Smith 2022).

**Figure 4.**
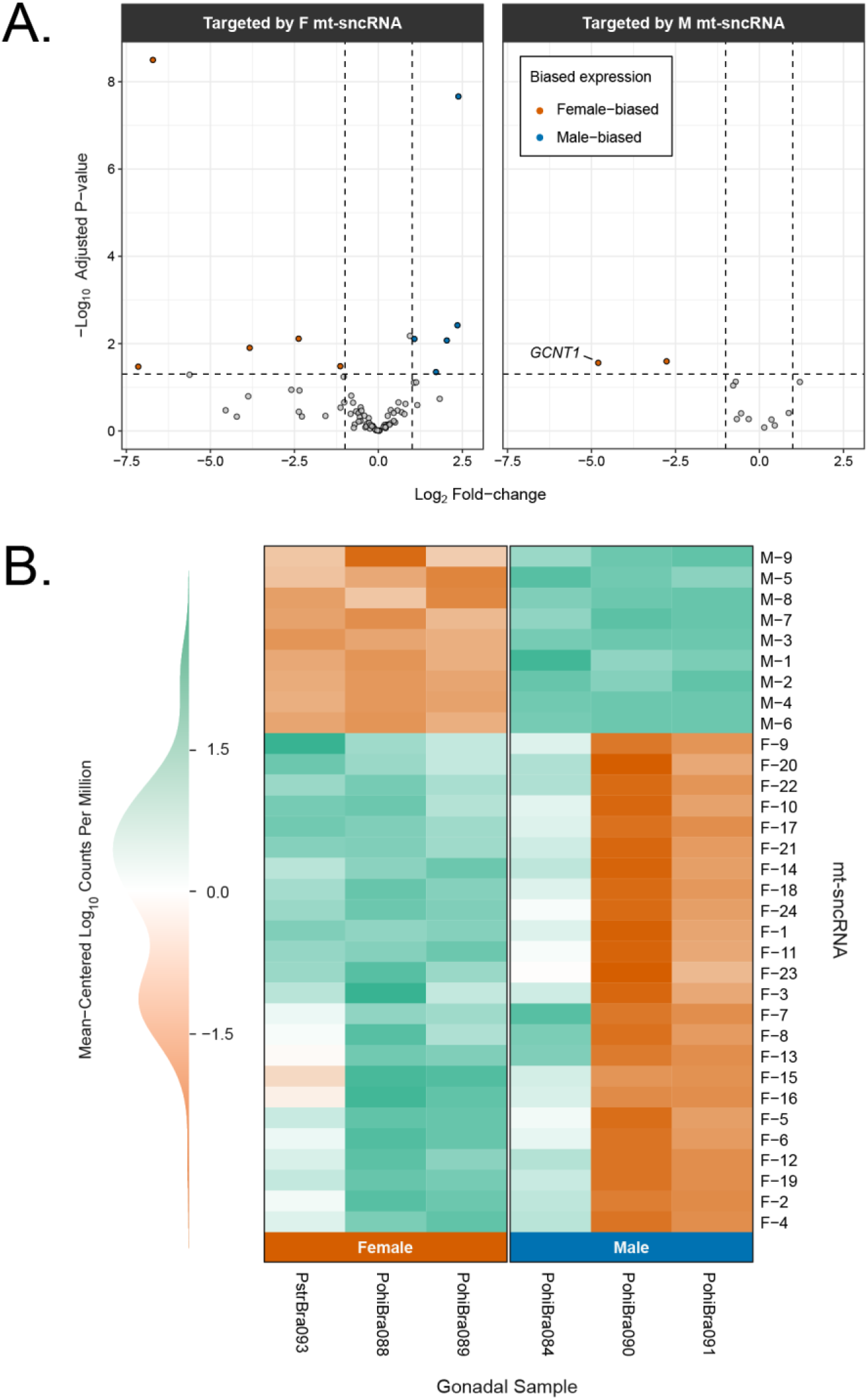
A) Volcano plot summarizing gene expression of genes targeted by female (left) and male mt-sncRNAs (right). Orange dots represent genes found to have significant female-biased expression and blue those with significant male-biased expression. All targets with significant differences in expression were hypothetical other than *GCNT1*. B) Heat map visualizing mt-sncRNAs expression in female (left) and male (right) gonadal tissue. Green coloration represents higher expression and orange represents lower or no exp ression.

We would generally expect nuclear targets of mt-sncRNAs to show sex-biased expression, with M mt-sncRNA targets most likely showing increased expression in females and vice versa for F mt-sncRNAs. However, we found limited expression differences in F and M mt-sncRNA nuclear gene targets (Fig. 4). Of the 96 mt-sncRNA targets identified, only 13 were sex-specific DEGs. Although we would expect all “true” mt-sncRNAs to lead to differential expression, a chi-squared test did support the hypothesis that mt-sncRNA targets were significantly enriched for DEGs (14% vs 6.6%, p = 0.006). Of the five F mt-sncRNA targets that follow the expected pattern of upregulation in males, the most extreme was ∼5-fold higher expression in male gonads (*Transient receptor potential melastatin 2* [*TRPM2*]). We also had seven F mt-sncRNA targets that showed female upregulation, with the most extreme having ∼141-fold higher expression in female gonads (*Endonuclease Domain Containing 1*). On the other hand, both M mt-sncRNA targets with differential expression were in the predicted direction and we choose to focus on the one with the most extreme sex-specific expression and also the only one that was annotated: a M mt-sncRNA located at the 5’ end of the M-ORF, which was predicted to target *Glucosaminyl (N-Acetyl) Transferase 1* (*GCNT1*), which had ∼28-fold higher expression in female gonads when compared to male gonads (log_2_-fold change = 4.8; p = 0.03) and in the top 5% of hub genes (Fig. 4; File S5). No M mt-sncRNA targets were upregulated in male gonads.

### F-ORF may interact with GCNT1 to facilitate female development

Because *GCNT1* showed the predicted patterns of a nuclear target gene under the CMS hypothesis and could therefore potentially play a role in sex differentiation of bivalves, we chose to further explore its potential interacting genes and proteins. The *GCNT1* gene network included 18 genes, most of which were *mucin* (*MUC*) and *N-acetylgalactosaminyltransferase* (*GALNT*) genes (Fig. 5). Many genes in the network were upregulated in females, including *GCNT1*, *MUC1* (log_2_-fold change = 2.10), *MUC4* (log_2_-fold change = 4.43), a duplicate of *MUC3A* (*MUC3A*_2; log_2_-fold change = 7), and a hypothetical gene showing homology to *MUC13* (*MUC13*-like; log_2_-fold change = 6.75) (Fig. 5). We found no acceptable support for PPIs between the GCNT1 protein and any of the mitochondrial proteins (*i.e.,* ATP8, F-ORF, and M-ORF), with the strongest interaction being between F-ORF and GCNT1 (Fig. S1). The potential F-ORF and GCNT1 PPI had a larger protein docking score (0.19) when compared to ATP8 (0.03) and the M-ORF (0.02). This is consistent with expectations from the CMS hypothesis but is limited evidence that these proteins interact.

**Figure 5.**
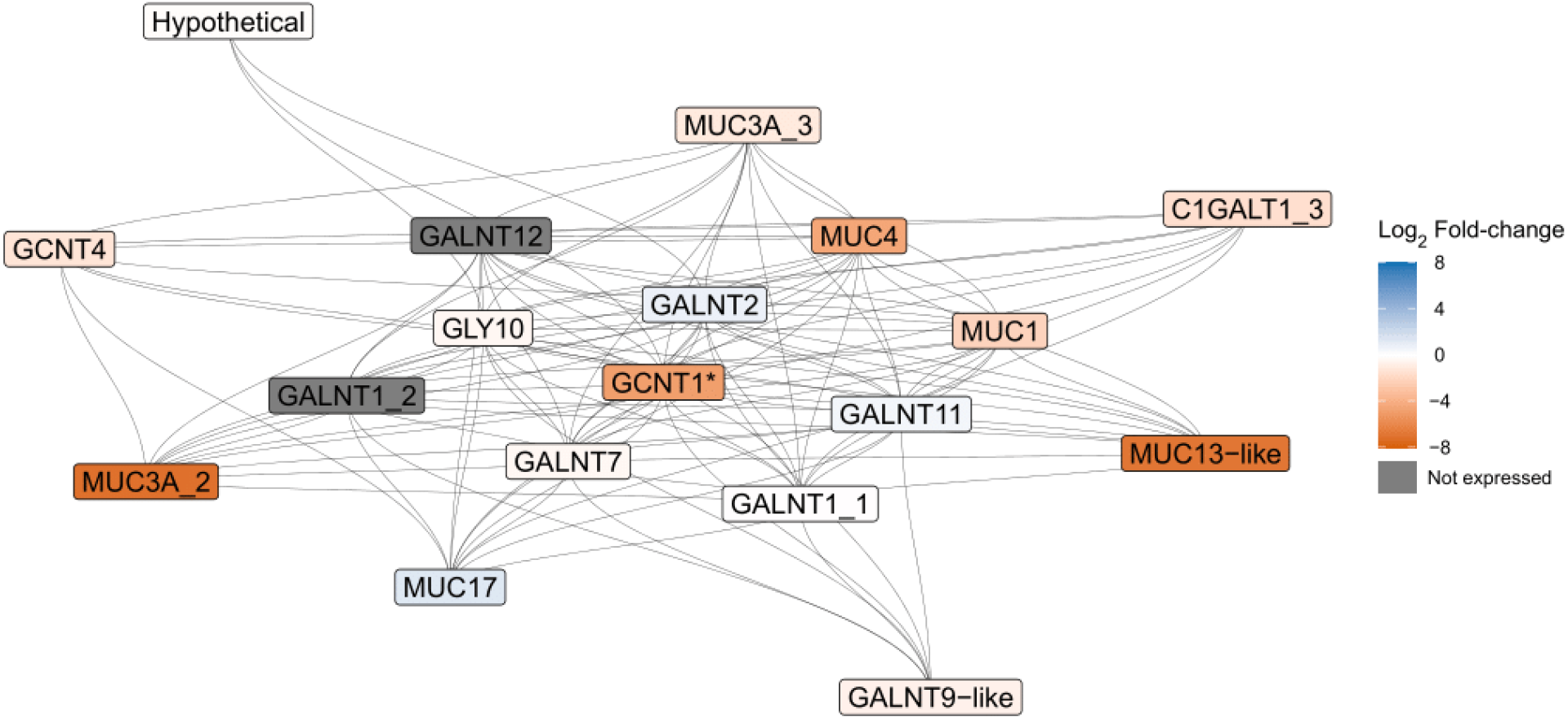
*GCNT1* gene network as predicted by STRING based on the *Potamilus streckersoni* proteome. Gray lines represent known interactions among genes based on database and experimental evidence. Genes are colored based on expression profiles between female (orange) and male (blue) gonadal tissue with color intensity representing differential gene expression between female and male gonadal tissue. Dark gray coloration represents genes with no expression. Asterisks indicated genes with significant differences in expression (p-adj < 0.05). Protein names with the suffix ‘-like’ are hypothetical and annotated as their best KEGG-hit when applicable. Protein names with an underscore represent protein products from genes with multiple annotated copies.

## Discussion

### Male mt-sncRNAs may inhibit mitophagy and contribute to sexual development in DUI species

In the CMS hypothesis presented by Breton et al. (2022), the F-ORF gene originated to act as a “feminizer” in an evolutionary transition from hermaphroditism to gynodioecy (*i.e.*, consisting of hermaphrodites and females). Subsequently, the M-ORF is hypothesized to have originated during a transition from gynodioecy to dioecy (*i.e.*, consisting of females and males) by counteracting feminization and leading to the origin of M mtDNA. While this hypothesis provides an explanation for the origin of DUI, empirical data has not supported that the M-ORF is responsible for counteracting the F-ORF or if other elements encoded within M mtDNA contribute to sex determination or sexual development. Our investigation found highly enriched mt-sncRNAs transcribed from M mtDNA that may play such a role (Fig. 4). Most interestingly, we validated a M mt-sncRNA at the 5’ end of the M-ORF that targeted *GCNT1*. This gene is in the mucin-type *O*-glycosylation pathway, which has a conserved role in eukaryotic development (reviewed by Tran and Ten Hagen 2013). Our RNA-seq data found *GCNT1* is differentially expressed and highly upregulated in females. This supports the hypothesis that M mtDNA is interacting with the nuclear genome through RNAi-mediated gene silencing and provides a plausible explanation for the evolutionary conservation of M-ORF across freshwater mussels.

The exact role that the knockdown of *GCNT1* plays in inhibiting degradation of M mtDNA and female developmental pathways remains uncertain, but our findings suggest two hypothetical functions. First, *GALNT* genes in the *GCNT1* gene network have been hypothesized to trigger female development in scallops based on comparative transcriptomics (Zhou et al. 2019), which supports the hypothesis that the M mt-sncRNA could inhibit female developmental pathways in freshwater mussels if it is evolutionarily conserved across bivalves. This hypothesis is further supported from findings in mice, where *GCNT1* has been demonstrated to be upregulated following the knockout of *SOX8* (Singh et al. 2009). These results suggest a non-altered male development pathway keeps *GCNT1* at low levels and provide further support for a role of M mt-sncRNAs in sexual development of bivalves through the inhibition of female developmental pathways.

Genes in the *GCNT1* gene network have also been demonstrated to play a role in mitophagy. The gene *MUC1* is localized to the mitochondrial membrane to increase mitophagy of mitochondria with decreased membrane potential (Li et al. 2022). Our results indicated that *MUC1* is somewhat upregulated in female gonads of *P. streckersoni* (log_2_-fold change of 2.1), which supports the hypothesis that upregulation of the *GCNT1* gene network in female gametes is involved in mitophagy of male-transmitted mitochondria and provides a plausible explanation for the M mt-sncRNA target. This hypothesis does require that male-transmitted mitochondria have a lower membrane potential than female-transmitted mitochondria. Decreased membrane potential has been demonstrated to trigger the degradation of paternal mitochondria post fertilization in multiple model species (Rojansky et al. 2016; Zhou et al. 2016). While sperm mitochondria in some marine DUI species have been demonstrated to exhibit high membrane potential (Milani and Ghiselli 2015), mitochondrial performance and function is certainly altered in DUI sperm vs. eggs (Bettinazzi et al. 2019; Bettinazzi et al. 2020) and M mtDNA OXPHOS genes have been suggested to be under relaxed selection (Maeda et al. 2021), suggesting that altered mitochondrial function in sperm could target M mtDNA for degradation. Studies of F vs. M mtDNA function have not been performed in freshwater mussels and are necessary to determine if male-transmitted mitochondria have decreased membrane potential compared to female-transmitted mitochondria, which would further support the role of the *GCNT1* gene network in degradation of M mtDNA during early development.

### No evidence of nuclear sex determination in freshwater mussels

Our findings did not show evidence of a nuclear SDR. This result is not surprising given bivalves lack heteromorphic sex chromosomes, and that SDRs on homomorphic sex chromosomes can be small. However, genome-wide heterozygosity was also unable to identify regions that were enriched for SDR-like alleles using similar methodologies as previous studies in bivalves (Han et al. 2022). Although we did not find evidence of nuclear genes involved in sex determination, we cannot reject that sex is determined by nuclear genes (Kenchington et al. 2009; Ghiselli et al. 2012; Milani et al. 2013; Zouros and Rodakis 2019; Zouros 2020), or directly by the male-transmitted mitochondria (Breton et al. 2011; Breton et al. 2022).

Under the hypothesis of nuclear sex determination, one or more loci could cause the retention or degradation of male-transmitted mitochondria in early development, which may or may not contribute to sexual development. This hypothesis does not require fixed differences between the sexes. We agree with previous hypotheses that nuclear encoded loci must contribute to the degradation or retention of male-transmitted mitochondria in early development, because while mitochondrial loci most likely contribute to more than just respiration, they must interact with nuclear gene products to perform any function (Rand 2001). We did observe high levels of genetic differentiation in numerous genes across the genome of *P. streckersoni* (1346 genes across 1/3 of all scaffolds) concomitant to differential expression of many of these genes, which is consistent with polygenic sex determination. However, this type of sex determination is hypothesized to be evolutionarily unstable (Rice 1986), and there are a only few hypothesized instances of this type of sex determining system across animals (Alexander et al. 2015; Roberts et al. 2016; Schartl et al. 2023). Future studies are necessary to test its potential presence in freshwater mussels.

### Hypothetical sexual development in freshwater mussels and future directions

We hypothesize that freshwater mussels either have a relatively small, cryptic ZW sex determining region that has remained undetected (as in other bivalves; Han et al. 2022), or have a polygenic sex determining system (Fig. 6). However, it is worth noting that nuclear factors may not be involved in sex determination. Crosses in several DUI lineages (*i.e.*, Mytilida, Unionida, Venerida) suggest that offspring sex is solely impacted by the mother and offspring sex ratios can vary from all female to all male progeny dependent on female brood (Saavedra et al. 1997; Kenchington et al. 2002; Ghiselli et al. 2011; Machordom et al. 2015). Although offspring sex being determined by the mother is consistent with ZW nuclear sex determination, extreme variation in sex ratios under natural conditions deviates from the expected sex ratios (50% of each sex). In *Mytilus*, mothers that give rise to all-male progeny are relatively rare (Saavedra et al. 1997; Kenchington et al. 2002), which is coincident in other groups with CMS and may suggest mtDNAs act as the primary sex determination signal. Evaluating each of these hypotheses in *P. streckersoni*, however, will require a more contiguous genome assembly, more thorough genome resequencing, and/or sequencing of parents and progeny from controlled crosses. Controlled crosses are feasible in captivity, but difficult for two reasons: 1) the freshwater mussel life cycle requires temporary larval encystment on vertebrate hosts to complete metamorphosis, and 2) most species reach sexual maturity at a relatively old age (1-2 years) compared to model systems (Barnhart et al. 2008; Haag 2012).

**Figure 6.**
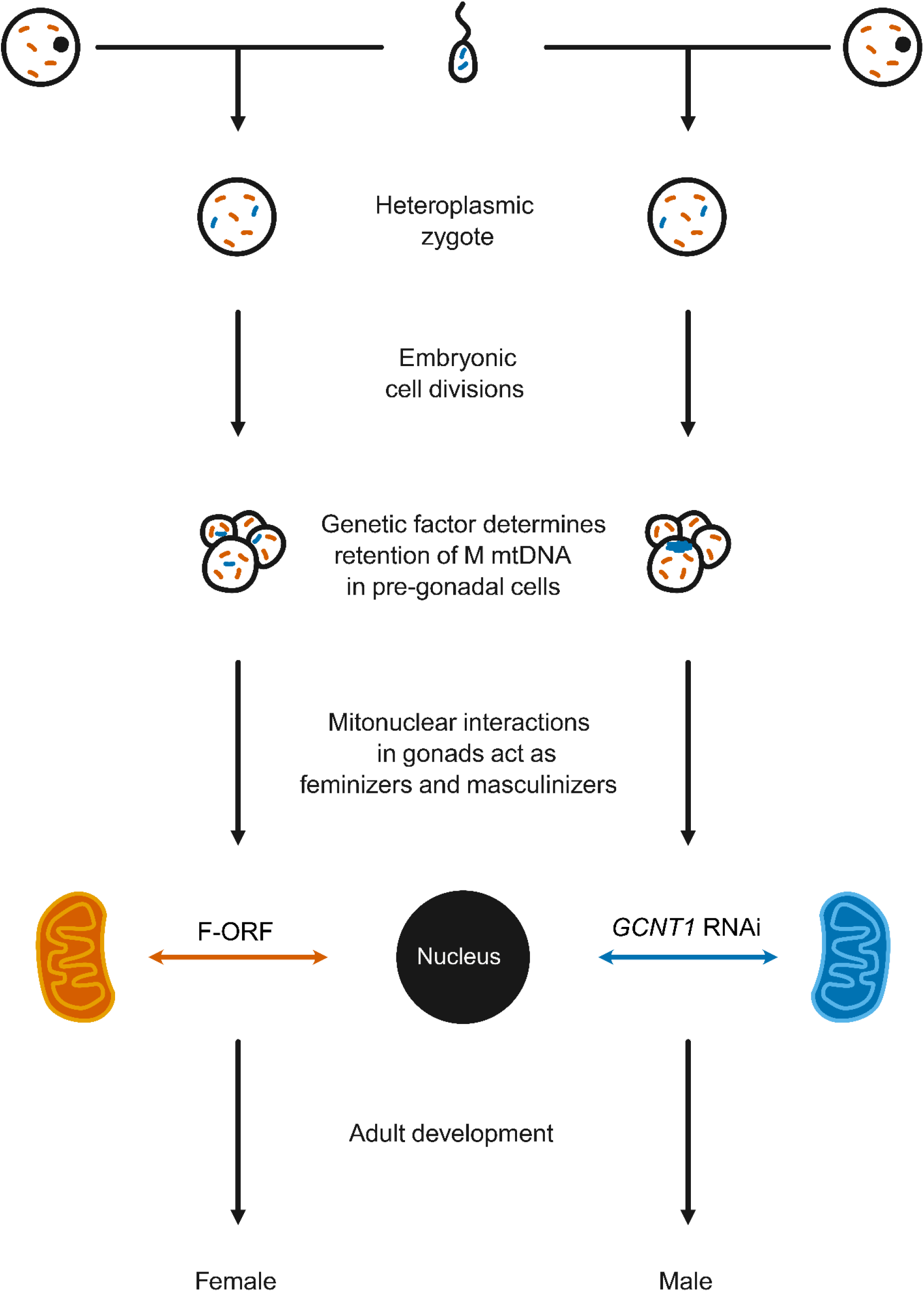
Hypothetical sexual developmental pathway in *Potamilus streckersoni*.

Despite not identifying a nuclear genetic sex determining region, our results do provide support for a direct role of M mt-sncRNAs acting as a “restorer” of maleness by directly inhibiting female developmental pathways through RNAi, with the mucin-type *O*-glycosylation pathway bring a prime target that may be important across bivalves. The knockdown of the pathway may inhibit female development or halt apoptosis of male-transmitted mitochondria in early development, but this remains speculative. Our hypothetical pathway is supported, at least in part, by empirical data and is consistent with previous hypotheses of sex determination in freshwater mussels (Breton et al. 2011; Breton et al. 2022). Although we did observe female upregulation of genes in the mucin-type *O*-glycosylation pathway (Fig. 5), we did not observe high genetic differentiation (F_ST_ ≥ 0.1) between the sexes in *GCNT1* or any other genes in the pathway. We also did not identify any ZW-like SNPs in the genes, which together with patterns of genetic differentiation, suggest that the pathway is epigenetically regulated. Future studies testing the regulatory roles of mt-sncRNAs, profiling gene and mt-sncRNAs expression during early development, and the use of mediated RNAi will be helpful in determining the role of mucin-type *O*-glycosylation pathway in female development. Further support could come from data on the *in vivo* activity of M mt-scnRNAs (as in Passamonti et al. 2020), their functional binding (Pozzi and Dowling 2022), their expression, and the expression of their targets during early development. Silencing of *GCNT1* through RNAi in female embryos will be helpful to determine if the mucin-type *O*-glycosylation pathway plays a role in female development. If the pathway is necessary for female development, silencing of that pathway should be strongly selected against in female embryos or lead to exclusively male progeny.

The role of the F-ORF or F mtDNA acting as a feminizer remains unclear. We were able to find limited support for the F-ORF protein interacting with the *GCNT1* (Fig. S1), but we cannot confirm that the two proteins interact. It is worth noting that the accuracy PPI prediction in non-model species is relatively poor (*e.g.*, Sledzieski et al. 2021). Future analytic advances or experimental datasets may provide empirical support for the F-ORF interacting with *GCNT1*. We were also able to validate 24 F mt-sncRNAs, but at this point we were unable to identify any targets that were supported by RNA-seq data to have a role in sexual development. It is worth noting that we were able to identify a putative F mt-sncRNA at the 5’ end of the F-ORF but could not validate a nuclear target at this time. Of the four genes targeted by F mt-sncRNAs that could be annotated (*i.e.*, *Fanconi Anemia Complementation Group E* [*FANCE*], *Krüppel associated box* [*KRAB*], *Nucleoporin 155*, *TRPM2*) and followed patterns expected under RNAi (*i.e.*, significant upregulation in males), we were unable to determine any biological functions potentially relevant to feminization. This is because the gene targets are broadly involved with complex gene pathways that provide multiple biological functions, albeit *FANCE*, *KRAB*, and *TRPM2* appear to have some level of contribution to mitophagy (Barde et al. 2013; Rodríguez and D’Andrea 2017; Kang et al. 2018). Future research will be necessary to infer the role of F mt-sncRNAs in female development, but we hypothesize most are involved in mitochondrial maintenance in gonadal and somatic cells.

## Data Availability Statement

All novel sequencing reads used in this study are available under BioProject PRJNA926666 on the NCBI SRA. Additional previously published reads used in the study are available under BioProject PRJNA681676. The updated genome annotation is available on NCBI under the WGS project accession JAEAOA01. Associated files and scripts used in this study are available in supplemental information and on GitHub (https://github.com/raquelmejiatrujillo/Mitonuclear_sex_determination_in_freshwater_mussels).

## Acknowledgements

The authors wish to thank the Cannatella, Havird, Hillis, Kirkpatrick, and Zamudio lab groups for useful comments on this manuscript. We also wish to thank staff at the Genomic Sequencing and Analysis Facility at the University of Texas at Austin, Center for Biomedical Research Support (RRID#: SCR_021713) and the Texas A&M AgriLife Genomics and Bioinformatics Service for assistance with data generation. This work was funded by the University Texas at Austin Stengl-Wyer Endowment and the National Institutes of Health (1R35GM142836).

