## Supplemental Figures, Files, and Tables for "Mitonuclear sex determination? Empirical evidence from bivalves": Supplementary Material.docx

**Supplemental Information**

***Supplemental Figures***

**Figure S1.** Protein-protein interaction (PPI) predictions for peptide sequences of: A) F-ORF and GCNT1, B) ATP8 and GCNT1, and C) M-ORF and GCNT1. Numbers on the x-axis label the two proteins and the y-axis reports the amino acid sequence length. Coloration corresponds to residue interaction predictions with lower values (blue) representing higher support for PPI.

***Supplemental File Descriptions***

**File S1.** Statistics from sliding 1kb windows as determined by PSASS. Columns are as follows: Contig – scaffold in *P. streckersoni* assembly, Position – start position of window on scaffold, Length – length of scaffold, Snps_females – female pool specific SNPs, Snps_males – male pool specific SNPs, Fst – F_st_ of window between males and feamles, Abs_depth_females – female pool read depth, Abs_depth_males – male pool read depth, Rel_depth_females – read depth of females relative to males, Rel_depth_males – read depth of males relative to females, and Depth_ratio – read depth ratio between females and males.

**File S2.** Statistics from sliding 1kb windows with an F_ST_ > 0.1 that encompassed a gene as determined by PSASS. Gene expression data is also reported as determined by DESeq2. Column names follow descriptions for files S1 and S3.

**File S3.** Gene expression data as determined by DESeq2. Column names are as follows: row – gene ID from *P. streckersoni* annotation, baseMean – mean base expression across all samples, log2FoldChange - log_2_FoldChange in expression between female and male gonads (negative female biased, positive male biased), lfcSE – log fold change standard error, stat – wald statistic, pvalue – uncorrected p-value, padj – p-value corrected for multiple comparisons, Target_annotation – annotated gene name (if applicable), Target_Contig – scaffold in *P. streckersoni* assembly that houses gene.

**File S4.** Modules selected by WGCNA to best explain expression profiles from RNA-seq data. Column names are as follows: FUN_ID – gene ID from *P. streckersoni* annotation, kTotal – connectivity of each gene to all other genes, kWithin – connectivity of each gene to all other genes within the same module, kOut – connectivity of each gene outside of its module relative to within its module, kDiff – connectivity of each gene outside of its module relative to within its module, module – module name, target_annotation – annotated gene name (if applicable), and target_contig – scaffold in *P. streckersoni* assembly that houses gene.

**File S5.** Genes with the highest intramodular connectivity as determined by WGCNA. Column names are identical to file S4.

**File S6.** The 246 pathways in the Hallmark and GO Biological Processes (GOBP) gene sets with significant enrichment for females or males. Column names are as follows: pathway – name of pathway, pval – uncorrected p-value, padj – p-value corrected for multiple comparisons, log2err – expected error for the standard deviation of the p-value logarithm, ES – enrichment score, NES – normalized enrichment score , size – number of genes in pathway used in calculations, and leadingEdge – genes used analysis of the pathway.

***Supplemental Table Descriptions***

**Table S1**. Details about specimens used for whole genome resequencing, including the sex of the individual, museum catalog number, number of read pairs post trimming, average genome coverage as calculated by SAMtools, and NCBI SRA accession number. Museum abbreviations are as follows: JBFMC – Joseph Britton Freshwater Mussel Collection and UF – Florida Museum.

**Table S2**. Details about specimens used for RNA-seq, including the sex of the individual, tissue type, museum catalog number, number of read pairs post trimming, and NCBI SRA accession number. All specimens are deposited in the Joseph Britton Freshwater Mussel Collection (JBFMC).

**Table S3.** Annotation completeness assessment from BUSCO scores using the metazoan and the molluscan lineages for the annotation presented by Smith (2021) and the updated annotation from this study.

**Table S4.** Details about specimens used for sncRNA-seq, including the sex of the individual, museum catalog number, number of reads post trimming, and NCBI SRA accession number. All specimens are deposited in the Joseph Britton Freshwater Mussel Collection (JBFMC).

**Table S5.** Female (F) and male (M) mitochondrial short non-coding RNAs (mt-sncRNAs) validated in this study. The mt-sncRNA sequence, its location, and predicted targets are reported. KEGG numbers (best hit with e-value < 1e^-5^) are provided for hypothetical genes, which are denoted in parentheses following the gene name. Genes named hypothetical had no acceptable hits. Expression data for each gene is provided (log_2_FoldChange) and the adjusted p-value from the differential expression analysis in DESeq2 (padj). Also, we provide the level of intramodular connectivity (kWithin) and the assigned module (module) from the WGCNA, with NAs representing genes that were not assigned to a module.
