## Supplementary figures and images for "Mitonuclear sex determination? Empirical evidence from bivalves"

### FigureS1.pdf

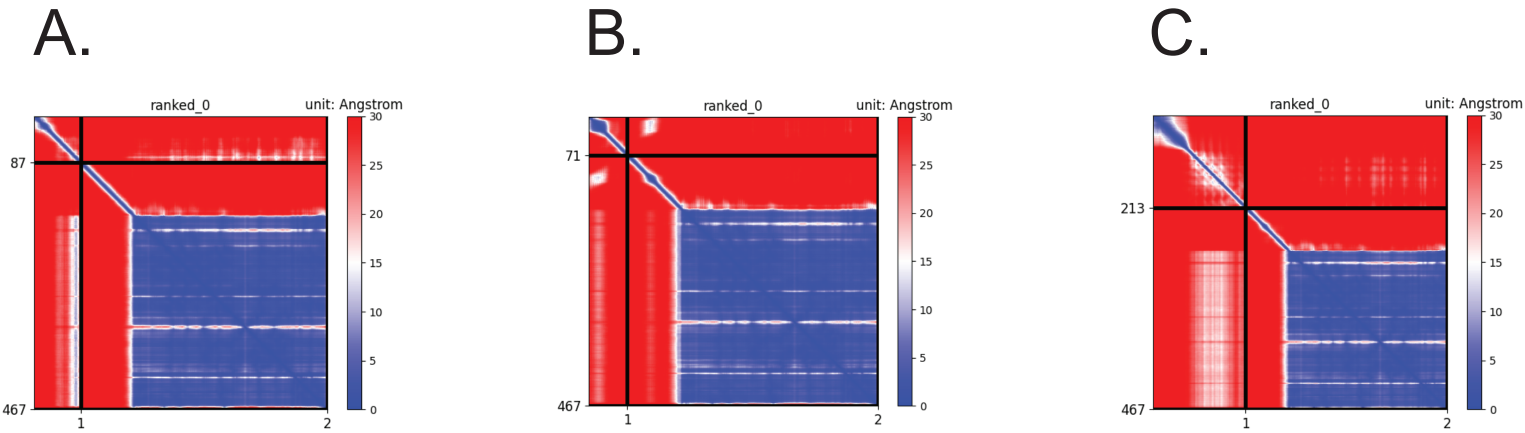
